# Dysregulated Proteins in Plasma Distinguishing Syndromic from Non-syndromic Heritable Thoracic Aortic Disease

**DOI:** 10.1101/2024.11.18.624213

**Authors:** Bjørn Edvard Seim, Yusuf Khan, Margrethe Flesvig Holt, Aleksandra Ratajska, Annika Michelsen, Monica Myklebust Ringseth, Runar Lundblad, Bente Halvorsen, Kirsten Krohg-Sørensen, Liv T. N. Osnes, Pål Aukrust, Benedicte Paus, John-Peder Escobar Kvitting, Thor Ueland

## Abstract

**BACKGROUND:** Thoracic aortic aneurysms (TAA) are often found in younger individuals and approximately 20% may be associated with heritable thoracic aortic disease (HTAD). There are some data on genomic biomarkers reflecting inflammation and extracellular matrix remodeling in HTAD. However, data that accurately reflect the corresponding protein changes are scarce. Our aim was to quantify proteins by using targeted proteomics in HTAD patients versus healthy controls (HC).

**METHODS:** Patients with Loeys-Dietz syndrome (LDS, *n* = 8), Marfan syndrome (MFS, *n* = 11), and Aortic aneurysm, familial thoracic 6 syndrome (AAT6, *n* = 7) were recruited. For comparison, blood samples were drawn from 16 healthy controls (HC). Plasma samples were analyzed by targeted proteome analysis of 276 proteins using immunoaffinity proteomics.

**RESULTS:** Seven proteins were elevated when comparing HTAD to HC, displaying two patterns: i) Oncostatin M (OSM) and Pentraxin-3 (PTX3) were significantly higher in both LDS and MFS, with comparable levels in AAT6, reflecting a generalized increase in HTAD. ii) increased levels of TNF Receptor Superfamily Member 9 (TNFRSF9), Granulysin (GNLY), Glycoprotein 1b-alpha (GP1BA), Vasorin (VASN) and CD5 were restricted to the LDS subgroup and correlated positively with Th17 and platelet counts.

**CONCLUSIONS:** The general increase in OSM and PTX3 in HTAD may reflect early but common mechanisms related to vascular inflammation while the distinct increase in TNFRSF9, GNLY, VASN, GP1BA and CD5 in LDS patients, may reflect involvement of Th17 inflammatory response mechanisms in the progression of TAA.

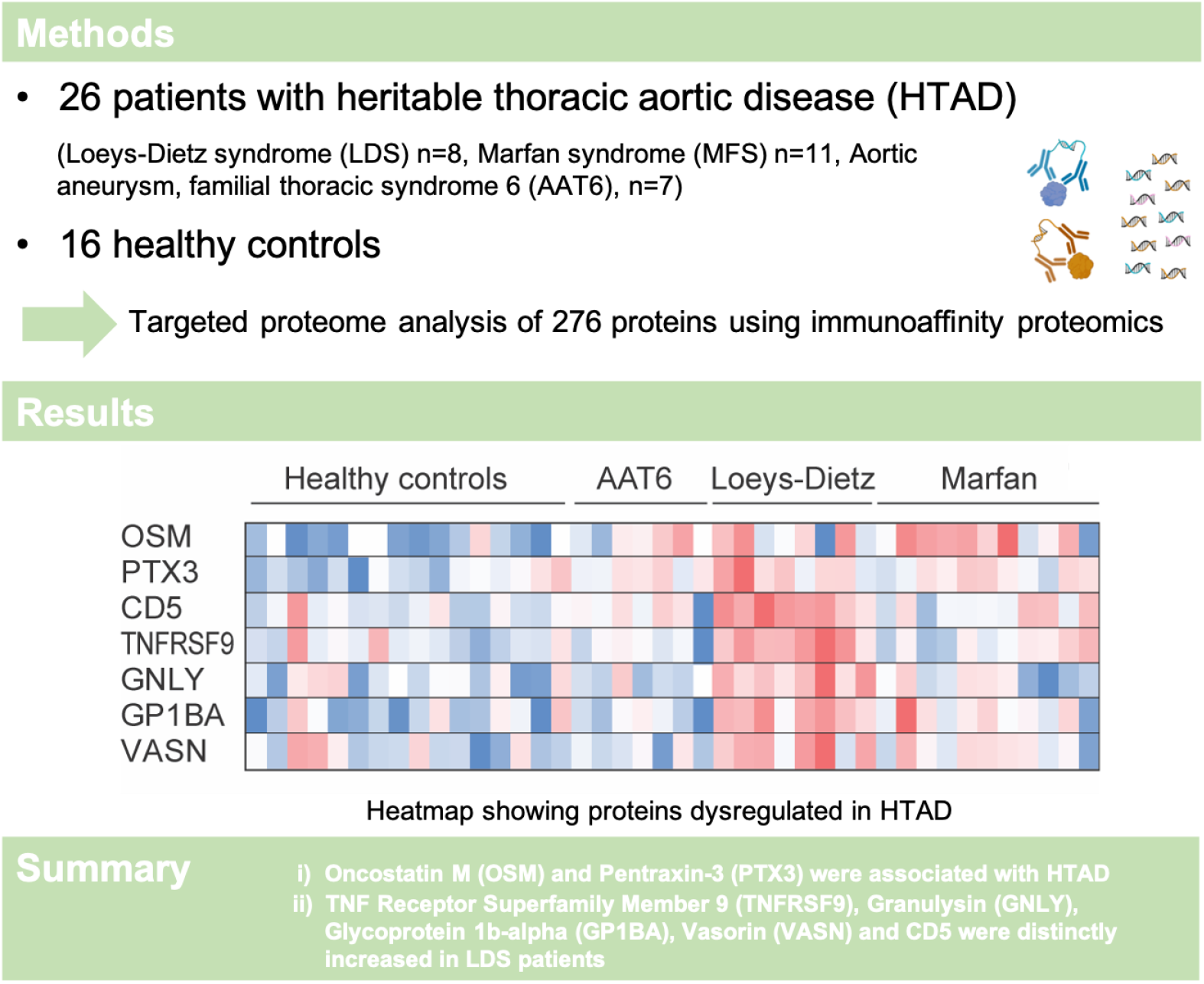

## INTRODUCTION

Whereas abdominal aortic aneurysms are associated with known risk factors for atherosclerosis (1), thoracic aortic aneurysms (TAA) are commonly found in younger patients and approximately 20 % may be associated with an underlying genetic disposition (2). Heritable thoracic aortic disease (HTAD) occurs as either syndromic, with extra-aortic features, or non-syndromic limited to the aorta. Both forms are frequently inherited in an autosomal dominant pattern. Genes affecting or interacting with transforming growth factor (TGF) signaling, genes related to the development and function of smooth muscle cells (SMC) and genes encoding the extracellular matrix (ECM) crosslinking enzymes have been associated with HTAD (2, 3). However, although genetic factors may point towards specific causal mechanisms, the understanding of the pathogenesis of HTAD is still incomplete. Due to its asymptomatic nature, there is also a need for non-invasive biomarkers and imaging tools to monitor HTAD.

Studies on genomic biomarkers that reflect inflammation or ECM remodeling in patients with HTAD are limited by the fact that they only provide indirect measurements of cellular states. However, this does not necessarily accurately reflect the corresponding protein changes (4). We have previously shown that plasma levels of matrix metalloproteinase (MMP)-9, a marker of ECM remodeling, and pentraxin (PTX) 3, a marker of vascular inflammation, are elevated in subgroups of patients with HTAD (5). Targeted protein screening offers a unique way to investigate a selected part of the proteome in the search for new biomarkers and mediators. To search for relevant biomarkers as well as to further elucidate the pathogenesis of HTAD, we therefore used a targeted proteome approach to quantify proteins associated with subgroups of HTAD including both syndromic (*i*.*e*. Loeys-Dietz syndrome [LDS] and Marfan syndrome [MFS]) and non-syndromic (*i*.*e*., Aortic aneurysm, familial thoracic 6 [AAT6]) forms. The protein levels were compared with levels in healthy controls as well as related to clinical characteristics of the different HTAD subgroups.

## MATERIALS AND METHODS

### Patients and healthy controls

The study was approved by the Regional Committee for Medical and Health Research Ethics in South-East Norway (REC no. 2018/732). All participants gave written informed broad consent. Patients were recruited at the multidisciplinary outpatient clinic for patients with vascular connective tissue diseases at the Department of Cardiothoracic Surgery, Oslo University Hospital (OUH), Rikshospitalet, Oslo, Norway. The patients were allocated into three groups of HTAD: LDS (n = 8), MFS (n = 11) and AAT6 (n = 7) as described previously (5). For comparison, blood samples were drawn from 16 healthy controls with no current or chronic diseases and no regular medications (Table 1). All patients were genetically characterized and had undergone exome or genome based sequencing analysis at the Department of Medical Genetics, OUH. Pathogenicity of variants were assessed by the American College of Medical Genetics and Genomics/Association for Molecular Pathology (ACMG/AMP) criteria, and all patients had a sequence variant that was assessed as pathogenic or likely pathogenic (6). All MFS patients had a pathogenic variant in *FBN1*, all LDS patients had a pathogenic variant in *TGF-βR1, TGF-βR2, SMAD3* or *TGF-β2*, and all AAT6 patients had a pathogenic variant in *ACTA2*.

**Table 1.**
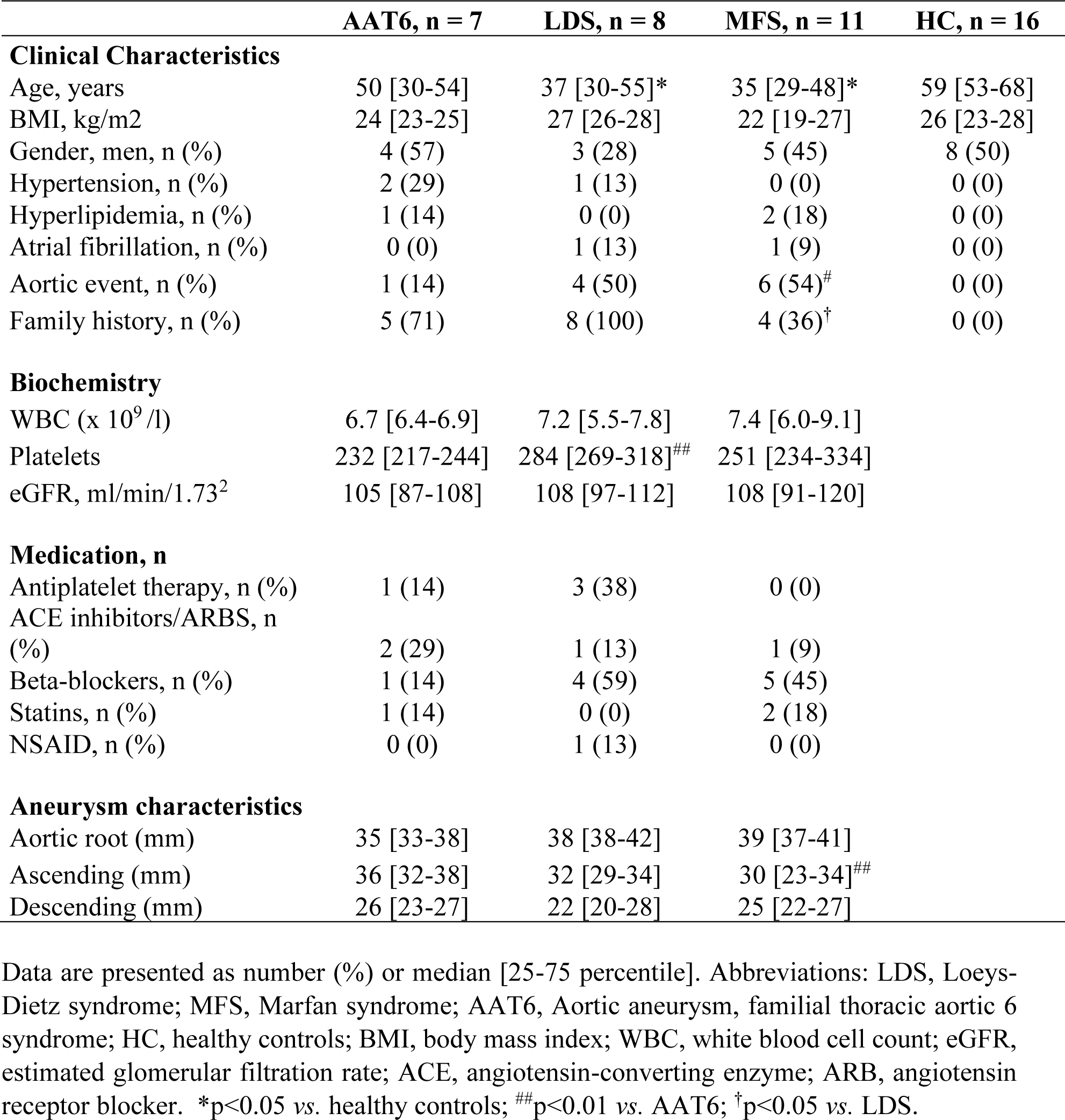
Demographic characteristics.

|  | **AAT6, n = 7** | **LDS, n = 8** | **MFS, n = 11** | **HC, n = 16** |
| --- | --- | --- | --- | --- |
| **Clinical Characteristics** |  |  |  |  |
| Age, years | 50 [30-54] | 37 [30-55]* | 35 [29-48]* | 59 [53-68] |
| BMI, kg/m<sup>2</sup> | 24 [23-25] | 27 [26-28] | 22 [19-27] | 26 [23-28] |
| Gender, men, n (%) | 4 (57) | 3 (28) | 5 (45) | 8 (50) |
| Hypertension, n (%) | 2 (29) | 1 (13) | 0 (0) | 0 (0) |
| Hyperlipidemia, n (%) | 1 (14) | 0 (0) | 2 (18) | 0 (0) |
| Atrial fibrillation, n (%) | 0 (0) | 1 (13) | 1 (9) | 0 (0) |
| Aortic event, n (%) | 1 (14) | 4 (50) | 6 (54)<sup>#</sup> | 0 (0) |
| Family history, n (%) | 5 (71) | 8 (100) | 4 (36)<sup>†</sup> | 0 (0) |
| **Biochemistry** |  |  |  |  |
| WBC (x 10<sup>9</sup>/l) | 6.7 [6.4-6.9] | 7.2 [5.5-7.8] | 7.4 [6.0-9.1] |  |
| Platelets | 232 [217-244] | 284 [269-318]<sup>##</sup> | 251 [234-334] |  |
| eGFR, ml/min/1.73<sup>2</sup> | 105 [87-108] | 108 [97-112] | 108 [91-120] |  |
| **Medication, n** |  |  |  |  |
| Antiplatelet therapy, n (%) | 1 (14) | 3 (38) | 0 (0) |  |
| ACE inhibitors/ARBS, n (%) | 2 (29) | 1 (13) | 1 (9) |  |
| Beta-blockers, n (%) | 1 (14) | 4 (59) | 5 (45) |  |
| Statins, n (%) | 1 (14) | 0 (0) | 2 (18) |  |
| NSAID, n (%) | 0 (0) | 1 (13) | 0 (0) |  |
| **Aneurysm characteristics** |  |  |  |  |
| Aortic root (mm) | 35 [33-38] | 38 [38-42] | 39 [37-41] |  |
| Ascending (mm) | 36 [32-38] | 32 [29-34] | 30 [23-34]<sup>##</sup> |  |
| Descending (mm) | 26 [23-27] | 22 [20-28] | 25 [22-27] |  |

Data are presented as number (%) or median [25-75 percentile]. Abbreviations: LDS, Loeys-Dietz syndrome; MFS, Marfan syndrome; AAT6, Aortic aneurysm, familial thoracic aortic 6 syndrome; HC, healthy controls; BMI, body mass index; WBC, white blood cell count; eGFR, estimated glomerular filtration rate; ACE, angiotensin-converting enzyme; ARB, angiotensin receptor blocker. \*p<0.05 vs. healthy controls; <sup>##</sup>p<0.01 vs. AAT6; <sup>†</sup>p<0.05 vs. LDS.

### Plasma sample collection

We collected peripheral venous blood from each patient into sterile tubes containing EDTA as an anticoagulant. The tubes were placed on melting ice, centrifuged within 30 minutes at 2000*g* for 20 minutes to obtain platelet-poor plasma, aliquoted and frozen in a -80 °C refrigerator for future use. The samples were thawed a maximum of three times.

### Targeted proteome analyses

Plasma samples from the HTAD groups (n = 26) and healthy controls (n = 16) were analysed using immunoaffinity proteomics. The Olink Target 96 panels (Olink Proteomics AB, Uppsala, Sweden) Cardiovascular II, Cardiometabolic, and Inflammation were assessed following the manufacturer’s guidelines at Proteomics Core Facility, Department of Immunology, University of Oslo/OUH. Subsequently, the Ct data were standardized and normalized by a set of internal and external quality controls. Finally, the raw data are log2 transformed and returned as Normalized Protein eXpression (NPX) values (www.olink.com).

### Statistical Methods

Demographics between the different diagnostic groups versus healthy controls were compared using Kruskal-Wallis *a priori*, and if significant, differences between groups were compared by the Mann-Whitney U-test. Categorical variables were compared using a chi square test. We performed proteomic data analysis using R (version 4.4.0) and R Studio (version 2023.03.0). We conducted Anova test using the OlinkAnalyze R package. All p-values were adjusted for multiple testing using the Benjamini-Hochberg procedure controlling the false discovery rate (FDR) at 5%. We used multiple R packages for data visualization. Significant markers from the Olink discovery were assessed in relation to cellular indices including flow cytometry, cell counts (white blood count and platelets), soluble markers of leukocyte activation as well as clinical features using spearman correlation or Mann-Whitney U-test for categorical variables. A two-sided p < 0.05 was considered significant.

## RESULTS

The demographic characteristics of the three HTAD groups, i.e., LDS, MFS and AAT6 as well as healthy controls are shown in Table 1. The patients had no known cancers or inflammatory diseases in addition to their HTAD. Whereas there were no differences in gender between controls and the different HTAD subgroups, the LDS and MFS group were significantly younger than controls. Six patients (4 MFS, 2 LDS) were operated for thoracic aorta dissection prior to inclusion in the study and 5 after (2 MFS, 2 LDS, 1 AAT6) giving a total of 11 aortic events.

### Proteomic analysis

Of 276 proteins assessed across three Olink panels, 7 proteins (oncostatin M [OSM], PTX3, TNF Receptor Superfamily Member 9 [TNFRSF9], granulysin [GNLY], Glycoprotein 1b-alpha [GP1BA], vasorin [VASN] and CD5) were significantly regulated between groups in age, sex- and full FDR-adjusted analysis (see Table 2 for a brief description). Figure 1A shows a heatmap summarizing the regulation of these 7 proteins. As depicted in Figure 1B, whereas OSM and PTX3 levels were higher in both LDS and MFS patients as compared with healthy controls, the elevation of TNFRSF9, GNLY, VASN, GP1BA and CD5 as compared with healthy controls were restricted to the LDS group. The levels of these markers in the LDS patients were in general higher than in both the MFS and AAT6 group. In fact, none of the 7 proteins were significantly elevated in the AAT6 group as compared with controls (Figure 1B). Thus, assessed by ROC analysis, this gave an area under the curve (AUC) of 0.89 for OSM and 0.76 for PTX for discriminating syndromic HTAD (i.e., LDS and MFS) from HC and an AUC between 0.84 and 0.97 for discriminating LDS from the other HTAD groups and HC with TNFRSF9 having the highest AUC.

**Table 2.** Brief description of proteins increased in heritable thoracic aortic disease.

| **Proteins** | **Descriptions and functions** |
| --- | --- |
| OSM, Oncostatin M | A pleiotropic, pro-inflammatory cytokine, expressed on endothelial cells, belonging to the IL-6 group of cytokines with a role in extracellular matrix remodeling and plaque vulnerability (33) |
| GNLY, Granulysin | An antimicrobial peptide released by killer lymphocytes in cytotoxic granules. Pore-forming peptide formed by cytotoxic immune cells (34). |
| CD5 | Cluster of differentiation (CD) expressed on the surface of most T cells and is upregulated on T cells upon strong activation and may amplify Th17 responses (24). |
| GP1BA, Glycoprotein 1ba | Also known as (CD42) function as a membrane receptor on platelet surface for von Willebrand factor allowing platelet adhesion (35). |
| TNFRSF9, 4-1BB, CD137 | TNF superfamily. Clonal expansion, survival and development of CD8+ T – cells. Lymphocyte activation (36). |
| PTX3, Pentraxin-3 | Produced and released by vascular cells in response to primary inflammatory signals. Acute and chronic inflammation (elevated in Loeys-Dietz syndrome) (37). |
| VASN, Vasorin | A transforming growth factor beta-binding protein expressed in vascular smooth muscle cells. Negative regulation of TNF signaling pathway and blocking TGF-β activity (28). |

**Figure 1.**
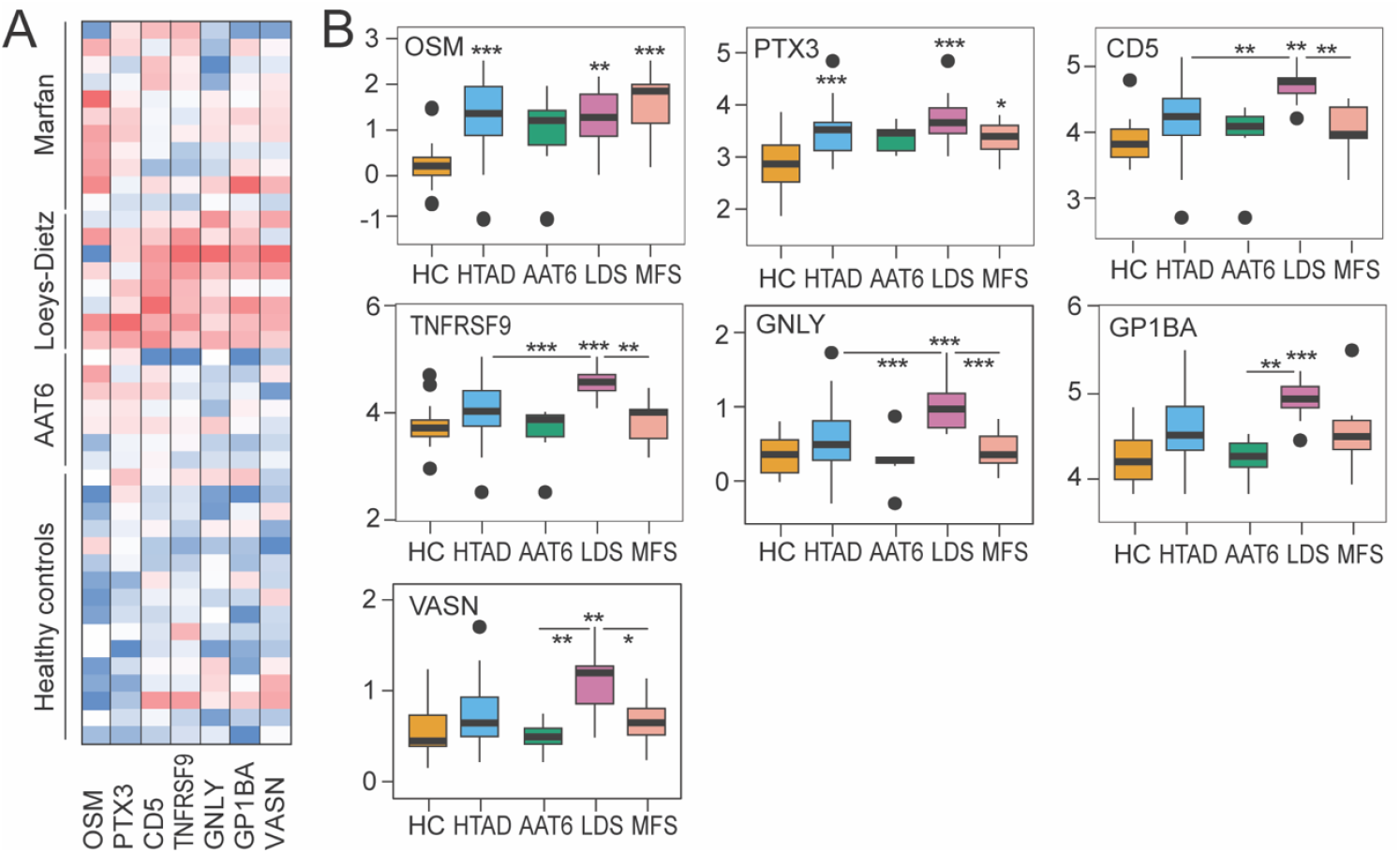
Dysregulated proteins in patients with heritable thoracic aortic disease (HTAD). A) Heatmap showing proteins dysregulated in HTAD and diagnostic subgroups using 3 Olink panels; OSM, oncostatin M; PTX3, pentraxin-3; TNFRSF9/4-1BB, TNF Receptor Superfamily Member 9; GNLY, granulysin; GP1BA, Glycoprotein 1b-alpha; VASN, vasorin. B) Tukey plots of normalized protein expression levels in Healthy controls (HC), HTAD and diagnostic subgroups: AAT6, Aortic aneurysm, familial thoracic 6 syndrome; LDS, Loeys-Dietz syndrome; MFS, Marfan syndrome. *p<0.01, **p<0.01, ***p<0.001 *vs*. HC or between indicated groups. Black dots indicate outliers.

In contrast to the association with subgroups of HTAD, these proteins were not associate with co-morbidities or medications listed in Table 1.

### Potential sources of dysregulated proteins

To identify potential circulating sources and targets of the dysregulated proteins we next assessed associations with previously determined cellular indices including flow cytometry (5), cell counts (white blood cell count and platelet count), soluble markers of leukocyte activation (monocyte/macrophage, sCD163 (7); T cell, sCD25 (8); neutrophils, myeloperoxidase [MPO] (9) and measures of TAA).

As shown in the heatmap in Figure 2A, some distinct patterns emerged for markers increased in patients with LDS (i.e., CD5, TNFRSF8, GNLY, GP1BA and VASN). These associations are visualized in Figure 2B. First, these markers tended to correlate positively with CD3 T cell percentage (p<0.05 for CD5, GNLY, GP1BA; p<0.08 for TNFRSF9 and VASN) and in particular Th17 counts (p<0.002 for GNLY, GP1BA and VASN; p=0.024 for TNFRSF9). Second, GNLY, GP1BA and VASN correlated positively with platelet count (p=0.006 for GP1BA and p<0.05 for GNLY and VASN). Third, for the two markers increased in both the LDS and the MFS group, OSM levels correlated negatively with naive CD4 T+ counts (p=0.037) and positively with Th1 (p=0.023) and WBC (p=0.004). PTX3 levels were negatively correlated with Th2 counts (p=0.007) and positively with MPO levels (p=0.042).

**Figure 2.**
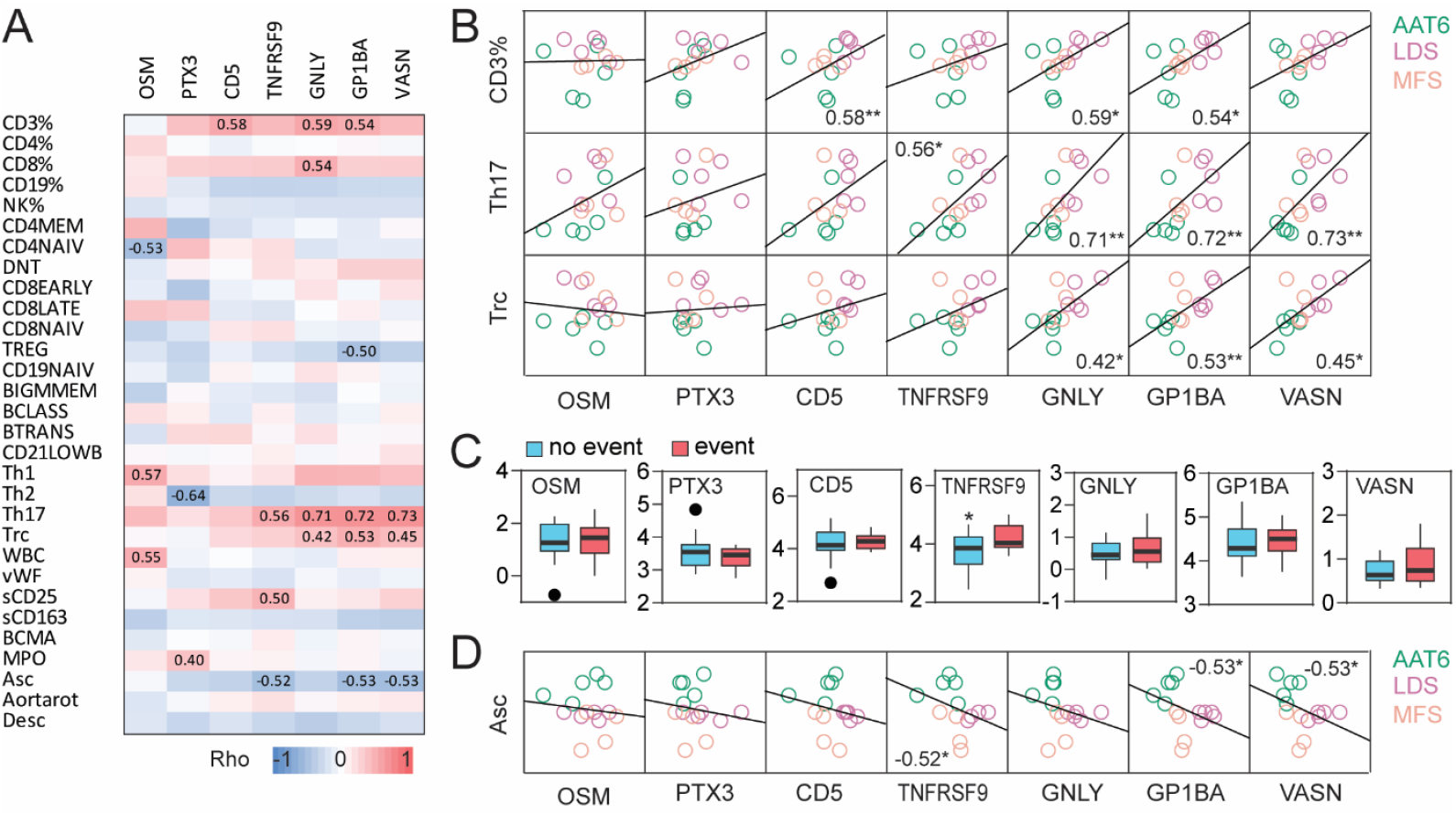
Potential sources of dysregulated proteins in heritable thoracic aortic disease (HTAD). **A)** Associations (spearman correlation) between dysregulated proteins (OSM, oncostatin M; PTX3, pentraxin-3; TNFRSF9/4-1BB, TNF Receptor Superfamily Member 9; GNLY, granulysin; GP1BA, Glycoprotein 1b-alpha; VASN, vasorin) in HTAD and previously determined cellular indices including flow cytometry, cell counts (WBC, white blood count; Trc, platelet count), soluble markers of leukocyte activation (monocyte/macrophage, sCD163; T cell, sCD25; neutrophils, MPO, myeloperoxidase) and measures of aortic aneurysm (Asc, diameter of ascending aorta; Desc, diameter of descending aorta) and cardiac functions (cTn, cardiac troponin; NT-proBNP). Correlation coefficients (Rho) are shown for significant correlations. **B)** Correlation plots between proteins and selected cellular indices. ^*^p<0.05, ^**^p<0.01. AAT6, Aortic aneurysm, familial thoracic 6 syndrome; LDS, Loeys-Dietz syndrome; MFS, Marfan syndrome. **C)** Tukey plots of normalized protein expression levels in HTAD according to aortic events (i.e. the need for operation). *p<0.05. Black dots indicate outliers. **D)** Correlation plots between proteins and diameter of the ascending aorta.

### Associations of biomarkers and clinical variables

As shown in Figure 2C, only TNFRSF9 levels were higher in patients with an aortic event (e.g., aortic dissection and operation). Moreover, TNFRSF9, GP1BA and VASN correlated negatively with the diameter of the ascending aorta (Figure 2D).

## DISCUSSION

By assessing 276 proteins using targeted proteomics we found seven proteins markedly elevated when comparing HTAD to HC. Notably, most of these proteins have previously not been related to HTAD. These proteins displayed two patterns of association with HTAD: i) OSM and PTX3 levels were higher in both LDS and MFS, with comparable levels in AAT6, although not statistical significant compared to HC, reflecting a generalized increase in HTAD. ii) Increased levels of five proteins (TNFRSF9, GNLY, VASN, GP1BA and CD5) were restricted to the LDS subgroup and correlated positively with Th17 and platelet count; potentially reflecting interactions between this T cell subset and platelets in the progression of LDS.

Vascular inflammation is an important initiating factor in the development of TAA, leading to remodeling of the aortic wall, characterized by accumulation and activation of immunocompetent cells (10). Our study found OSM and PTX3 to be elevated in the LDS and MFS subgroup compared with HC. We have previously shown elevated PTX3 in LDS (5) and with this study extend these findings by assessing associations with different cellular markers. Both OSM and PTX3 proteins are expressed in various organs and may promote local inflammation in relevant cells in HTAD such as endothelial cells, macrophages and SMC (11, 12). However, our finding that OSM correlated with WBC count, mainly reflecting neutrophil numbers, and PTX3 with MPO, a marker of neutrophil activation and oxidative stress (13) may suggest that activated neutrophils also contribute to systemic levels. Both OSM and PTX3 are stored in neutrophil granules and may be released upon activation, contributing to modulation of local inflammation (14, 15). Neutrophil infiltration has been noted in TAA (16), promoting formation of aortic aneurysms by secreting neutrophil extracellular traps and IL-6 (17). We hypothesize that increased OSM and PTX3 plasma levels may reflect early but common mechanisms related to interactions between immune cells and endothelial cells in the formation and progression TAA in syndromic HTAD.

The other pattern found in the proteomic analysis was the restricted elevation of TNFRSF9, GNLY, VASN, GP1BA and CD5 in LDS patients. These proteins also displayed a similar pattern of association with cellular markers, in particular with various T cell subsets, as reflected by the strong association Th17 cells. Th17 cells have been implicated in the progression of abdominal aortic aneurysms, promoting local inflammatory responses (18), while inhibiting IL-17A in these cells attenuates aneurysm development in experimental models (19). Although Th17 cells has been related to the progression of aortic dissection after thoracic endovascular aortic repair (20), data on Th17 cells in TAA are scarce or lacking.

The detected proteins were in general also associated with other T cell subsets, and notably, T cells were the largest cell population identified in aneurysmal human aortic tissue with a high expression of cytokine receptors including TNFRSF9 (21). GNLY is exclusively expressed by activated cytotoxic T lymphocytes and natural killer cells, acting as a chemoattractant for many cell types, promoting expression of proinflammatory cytokines such as IL -1 and IL-6 (22). CD5 is thought to be a marker of B cell subsets, but is also constitutively expressed on lymphocyte precursors and mature T cells (23), and notably, may amplify Th17 responses (24). In contrast to the T cell related proteins, GP1BA is a surface membrane protein of platelets. Non-laminar blood flow in aortic aneurysms may enhance platelet activation and aggregation, with release of membrane receptors and chemotactic cytokines, further promoting accumulation of inflammatory cells. With relevance to our results, activation of platelets have been associated with dilation of human thoracic ascending aortic aneurysm (25). In addition to enhancing secretion of inflammatory mediators and oxidative stress, infiltrating immune cells may also stimulate SMCs to produce various proteases leading to structural remodeling of the aortic wall (26). The increased levels of VASN, a transmembrane glycoprotein abundantly expressed on the surface of SMCs in the aorta (27) could reflect involvement of SMCs in LDS but could also more directly contribute to the LDS phenotype that are related to pathogenic variants of genes encoding components of TGFβ family. In fact, VASN inhibits TGF signaling (28), corresponding with the LDS phenotype (29), pointing to a new mechanism in the attenuated TGF responses in these patients.

We speculate that increased levels of these T cell related proteins may reflect interactions between T cell subsets and platelets in enhancing aneurysm development through effects on structural integrity and stability of the vessel wall (30). TNFRSF9 levels were higher in patients with an aortic event of relevance. The ligand for this receptor, TNFSF9, enhances the production of inflammatory cytokines driven by TLR signaling and promotes macrophage M1 polarization (30), an important feature in aortic aneurysm progression (31) associated with rupture (32, 33). To the best of our knowledge, this is the first report of a potential pathogenic role of TNFRSF9 in HTAD. Conversely, plasma levels of TNFRSF9, GP1BA and VASN were negatively correlated with diameters of the ascending aorta. However, diameter has been reported to be a poor predictor of risk (3) and the threshold for thoracic surgery is also lower in LDS compared to MFS and AAT6 with ACTA2 mutations (31). Indeed, the AAT6 patients in our study were older, had a larger diameter of the ascending aorta, but fewer aortic events reflecting the less aggressive nature of the aortic disease in the AAT6 group compared to syndromic HTAD such as LDS and MFS.

Five of the seven dysregulated proteins were found in the LDS subgroup of HTAD patients. This could reflect the clinical impression that the LDS patients have a more severe disease manifestation than MFS and AAT6 patients. Current guidelines for optimal timing of prophylactic surgery in HTAD depend largely on aortic size, family history and symptoms (32), despite the high risk of life-threatening aortic events even at smaller aortic dimensions. The markers displayed descent discriminatory properties in identifying HTAD and in particular LDS, which should be validated in larger independent cohorts. Furthermore, these proteins, the cells and pathways they reflect, could be modified by novel interventions. In particular, targeting Th17/IL-17A–related inflammatory responses may have merit as shown in experimental models of abdominal aortic aneurysms (19). Combining these biomarkers with advanced imaging studies quantifying for example wall shear stress could potentially lead to a more personalized follow-up and treatment algorithm of the individual HTAD patient.

This study is limited by being a single center study with a small patient cohort. Based on this, the results must be interpreted with caution and the identified proteins need to be validated in independent cohorts. The subgroup of FTAAD consisted exclusively of patients with the ACTA2 mutation (e.g. AAT6). ACTA2 mutations, however, comprise 12% -21% of the FTAAD population. Furthermore, we did not validate our proteomic findings with independent methods. However, we have found a similar association between PTX3 plasma levels HTAD previously and correlation with PTX3 levels from the O-link assay were strong (r=0.89). Moreover, we lack data on clinical manifestation beyond that of aortic dimensions in the syndromic TAA patients. Finally, our results are mostly based on associations, which not necessarily mean any causal relationship. In particular, although our data suggest association of some of the markers with certain leukocyte subsets, we lack data on intracellular levels of these molecules in the actual cells.

In conclusion, of the seven dysregulated proteins found in this study, elevated OSM and PTX3 were associated with HTAD in general, while TNFRSF9, GNLY, VASN, GP1BA and CD5 where distinctly increased in LDS patients, potentially reflecting the involvement of Th17 related mechanisms in the progression of TAA. Larger studies, are needed to clarify whether these proteins could represent diagnostic or prognostic biomarkers that can be the basis for a more individualized follow-up and treatment.

## Non-standard Abbreviations and Acronyms

CD5: Cluster of differentiation 5
ECM: Extracellular matrix
AAT6: Aortic aneurysm, familial thoracic 6 syndrome
GNLY: Granulysin
GP1BA: Glycoprotein 1b-alpha
HTAD: Heritable thoracic aortic disease
LDS: Loeys-Dietz syndrome
MFS: Marfan syndrome
MMP-9: Matrix metalloproteinase-9
MFO: Myeloperoxidase
OSM: Oncostatin M
PTX3: Pentraxin-3
TAA: Thoracic aortic aneurysms
TGF: Transforming growth factor
TNFRSF9: TNF receptor superfamily member 9
VASN: Vasorin

## Acknowledgements

Proteomic analyses were performed by the Proteomics Core Facility, Department of Immunology, University of Oslo/Oslo University Hospital, which is supported by the Core Facilities program of the South-Eastern Norway Regional Health Authority. This core facility is also a member of the National Network of Advanced Proteomics Infrastructure (NAPI), which is funded by the Research Council of Norway INFRASTRUKTUR-program (project number: 295910).

## HIGHLIGHTS

- Oncostatin M (OSM) and Pentraxin-3 (PTX3) were significantly higher in both Loyes-Dietz syndrome (LDS) and Marfan syndrome, with comparable levels in Aortic aneurysm, familial thoracic syndrome 6, reflecting a generalized vascular inflammation in heritable thoracic aortic disease.
- Increased levels of TNF Receptor Superfamily Member 9, Granulysin, Glycoprotein 1b-α, Vasorin and CD5 were restricted to the LDS subgroup and correlated positively with Th17 and platelet counts.
- Our finding could potentially reflect the involvement of Th17 and platelet related mechanisms in the progression of LDS

